# Involvement of non-LTR retrotransposons in mammal cancer incidence and ageing

**DOI:** 10.1101/2021.09.27.461867

**Authors:** Marco Ricci, Valentina Peona, Cristian Taccioli

## Abstract

The presence in nature of closely related species showing drastic differences in lifespan and cancer incidence has recently increased the interest of the scientific community on these topics. In particular, the adaptations and genomic characteristics underlying the evolution of cancer-resistant and long-lived species have recently focused on the presence of alterations in the number of non-coding RNAs, on epigenetic regulation and, finally, on the activity of transposable elements (TEs). In this study, we compared the content and dynamics of TE activity in the genomes of four rodent and six bat species exhibiting different lifespans and cancer susceptibility. Mouse, rat and guinea pig (short-lived and cancer-prone organisms) were compared with the naked mole rat (*Heterocephalus glaber*) which is the rodent with the longest lifespan. The long-lived and cancer-resistant bats of the genera *Myotis*, *Rhinolophus*, *Pteropus* and *Rousettus* were instead compared with the *Molossus*, which is instead a short-lived and cancer-resistant organism. Analyzing the patterns of recent accumulations of TEs in the genome in these species, we found a strong suppression or negative selection to accumulation, of non-LTR retrotransposons in long-lived and cancer-resistant organisms. On the other hand, all short-lived and cancer-prone species have shown recent accumulation of this class of TEs. Among bats, the *Molossus molossus* turned out to be a very particular species and, at the same time, an important model because, despite being susceptible to rapid ageing, it is resistant to cancer. In particular, we found that its genome has the highest density of SINE (non-LTR retrotransposons), but, on the other hand, a total lack of active LINE retrotransposons. Our hypothesis is that the lack of LINEs presumably makes the *Molossus* cancer resistant due to lack of retrotransposition but, at the same time, the high presence of SINE, may be related to their short life span due to “sterile inflammation” and high mutation load. We suggest that research on ageing and cancer evolution should put particular attention to the involvement of non-LTR retrotransposons in these phenomena.

## Background

Transposable elements (TEs) are repeated mobile elements present in almost all eukaryotes [1, 2]. They can accumulate in genomes and account for most of the organism genome size, for example, the human genome is >50% repetitive [3]. Many studies have shown that TEs are implicated in gene duplications, inversions, exon shuffling, gene expression regulation and may play an important role in the long-term evolution of organisms [4–12]. For example, the co-option of TE-related proteins gave rise to vertebrate acquired immune system and mammalian placenta [13–19]. Although some TE insertions have been co-opted by genomes, the overall TE activity can be disruptive especially at short timescale. Indeed, TEs can be responsible for several human diseases [7, 20, 21] like immunodeficiency [22], coagulation defects [23], cardiomyopathies [24], and muscular dystrophies [25, 26]. Most interestingly, the dysregulation of TEs in somatic cells may be also involved in the establishment, development and plasticity of cancer [27–32]. The dysregulated TE activity in cancer cells is so pervasive and accentuated that the methylation level of transposable elements is used as a biomarker for the malignancy of several types of tumours [33–37]. Given the manifold effects of TEs on health, it is reasonable to consider the TE activity a key factor able to influence lifespan differences between species [38]. The experimental design presented here is based on the comparison of the TE activity in genomes of mammals with different lifespans and cancer incidence. The TEs that cause mutations and genomic instability in the genomes are the ones currently active and able to move throughout the genome [38–41]. Since there are no transposition essays available for many organisms, we used the genetic divergence of the TE insertions (see **Methods**) from their consensus sequences as a proxy for their active or inactive state [12]. Among all the mammalian species for which genome assemblies are publicly available, we chose four species of Rodentia and six of Chiroptera that: 1) have a high-quality genome assembly (in order to maximize the quality and quantity of transposable elements assembled [1]); 2) belong to the same taxonomical order (to maximize their shared evolutionary history); and 3) have comparable body masses. In particular, the inclusion of body mass among the criteria of selection allowed us to work with species that have not evolved biological adaptations to counteract cancer incidence as a function of body size (Peto’s paradox) [42]. In fact, animals with a big body mass evolved mechanisms to contrast the development of tumours such as the control of the telomerase expression [43] or the expansion in copy number of the onco-suppressor TP53 in *Loxodonta africana* (African elephant) [44]. Small mammals have a smaller number of cells and generally shorter life cycle [45], therefore they are disentangled from the Peto’s paradox. For this reason, we selected mammals with a body mass lower than 2 kg to investigate the relationship between lifespan and transposable elements avoiding biases related to adaptations to large body masses. The species we used in our analyses represent both long and short lifespans (**Figure 1**) providing the opportunity to investigate the relationship between the TE activity and ageing. Here we test the hypothesis that the genomes of cancer-prone and short-lived species present a higher load of recently inserted TEs than cancer-resistant and long-lived species.

**Figure 1.**
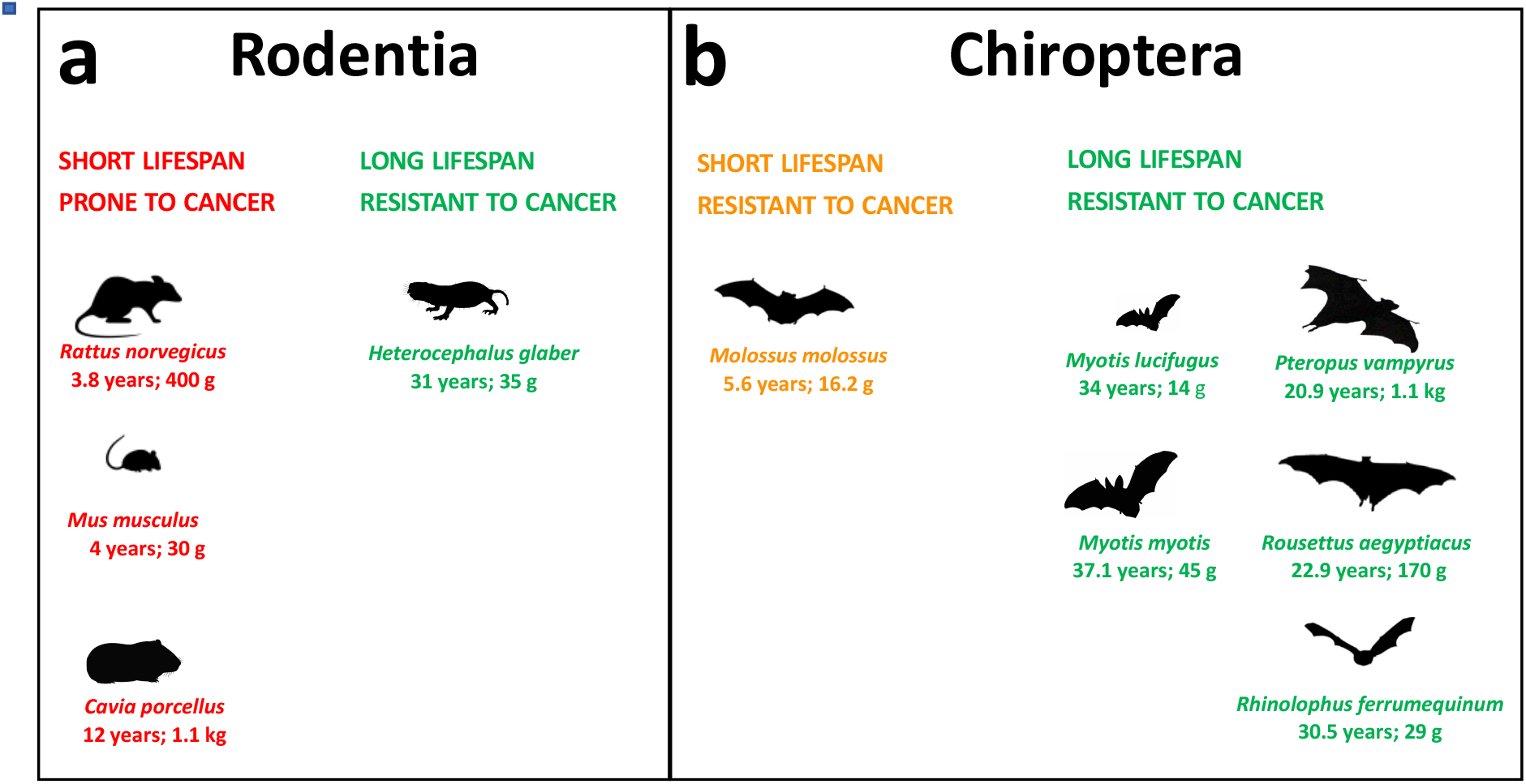
List of species analysed together with lifespan and body mass information. **a**) Rodents with short lifespan and high cancer incidence are indicated in red while species with long lifespan and low cancer incidence are indicated in green. **b**) Bats with short lifespan and low cancer incidence are indicated in orange while species with long lifespan and low cancer incidence are indicated in green. This information was retrieved from https://genomics.senescence.info/species/index.html and Speakman et al. [44].

## Results

In this study we investigated the possible link between TEs, cancer incidence, and ageing by focusing on the accumulation patterns of TEs in genomes belonging to species showing different lifespans and cancer incidence (**Figure 1**). First, we collected genome assemblies of similar high quality and generated a TE library for the species that previously lacked one to both maximise the presence of TEs in the assemblies and their correct annotation. In particular, we showed that the last genome assembly version of *Heterocephalus glaber* (naked mole rat) based on long reads, yielded a higher percentage of the genome annotated as TEs with respect to its first version based on short reads (**Figure 2**, **Table 1**). Second, we proceeded with the comparison of the TE accumulation profiles between short- and long-lived rodents and observed a reduced accumulation of non-LTR retrotransposons in the short-lived species (**Figure 3**). The same analysis in bats showed a high accumulation of class II transposons in all species (**Figure 4**, **Table 2**) but also highlighted a drop in the accumulation of non-LTR retrotransposons in the genomes of the long-lived bats (**Figure 4**). Finally, a detailed analysis of the most recent accumulation of non-LTR retrotransposons in rodents and bats showed a higher accumulation density in the species with the lowest lifespan (**Figure 5** and **6**).

**Table 1.** The assembly size and the percentage of genome annotated as transposable elements are shown for each rodent genome.

| Species | *H. glaber* | *C. porcellus* | *M. musculus* | *R. norvegicus* |
| --- | --- | --- | --- | --- |
| Assembly size (Gb) | 3.042 | 2.723 | 2.728 | 2.648 |
| Total TE content (%) | 38.16 | 36.01 | 41.86 | 41.76 |
| DNA transposons (%) | 2.3 | 1.7 | 1.1 | 1.1 |
| SINE (%) | 4.3 | 5.5 | 7.6 | 7.2 |
| LINE (%) | 24.78 | 21.4 | 19.2 | 20.9 |
| LTR (%) | 5.7 | 7.4 | 12.6 | 10.7 |
| Unknown (%) | 1.2 | 1.7 | 1.4 | 1.9 |

**Table 2.**
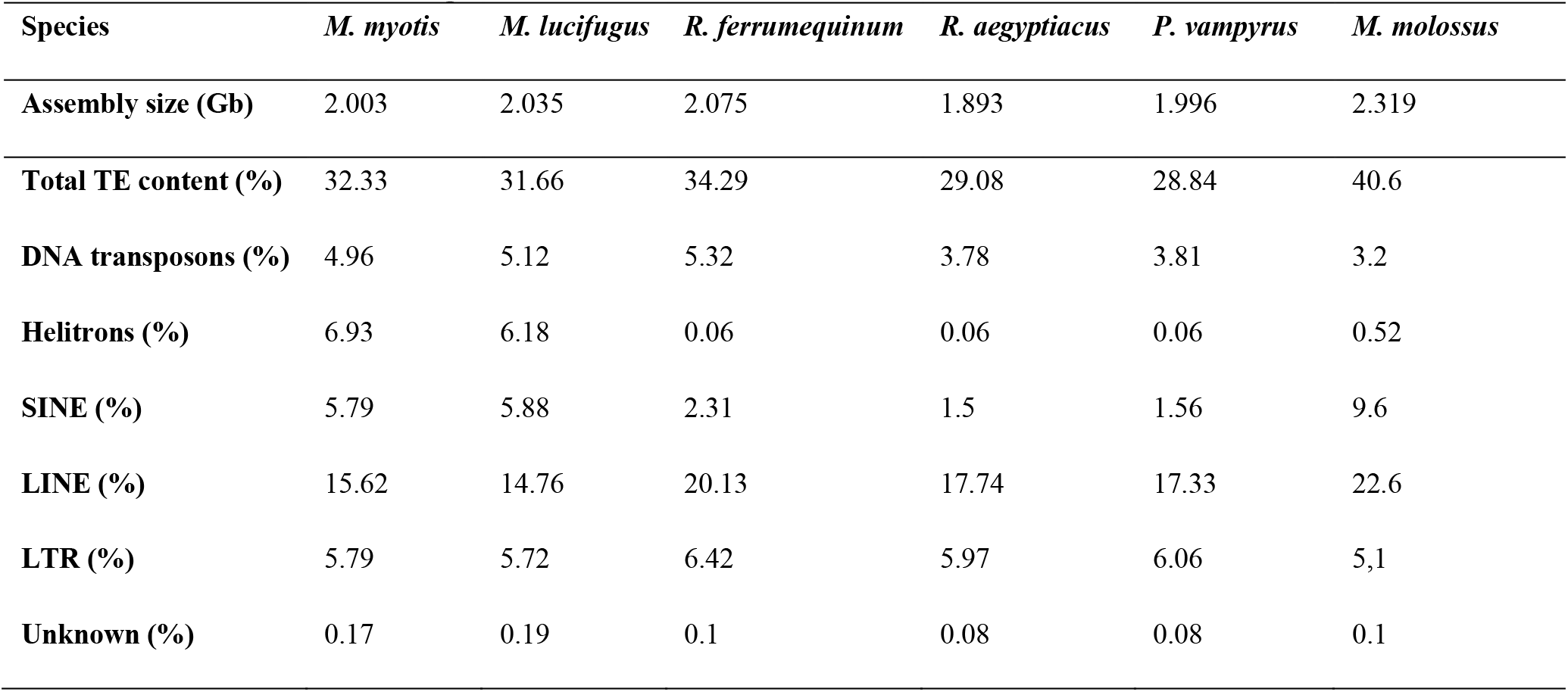
The assembly size and the percentage of genome annotated as transposable elements are shown for each bat genome.

| Species | *M. myotis* | *M. lucifugus* | *R. ferrumequinum* | *R. aegyptiacus* | *P. vampyrus* | *M. molossus* |
| --- | --- | --- | --- | --- | --- | --- |
| Assembly size (Gb) | 2.003 | 2.035 | 2.075 | 1.893 | 1.996 | 2.319 |
| Total TE content (%) | 32.33 | 31.66 | 34.29 | 29.08 | 28.84 | 40.6 |
| DNA transposons (%) | 4.96 | 5.12 | 5.32 | 3.78 | 3.81 | 3.2 |
| Helitrons (%) | 6.93 | 6.18 | 0.06 | 0.06 | 0.06 | 0.52 |
| SINE (%) | 5.79 | 5.88 | 2.31 | 1.5 | 1.56 | 9.6 |
| LINE (%) | 15.62 | 14.76 | 20.13 | 17.74 | 17.33 | 22.6 |
| LTR (%) | 5.79 | 5.72 | 6.42 | 5.97 | 6.06 | 5,1 |
| Unknown (%) | 0.17 | 0.19 | 0.1 | 0.08 | 0.08 | 0.1 |

**Figure 2.**
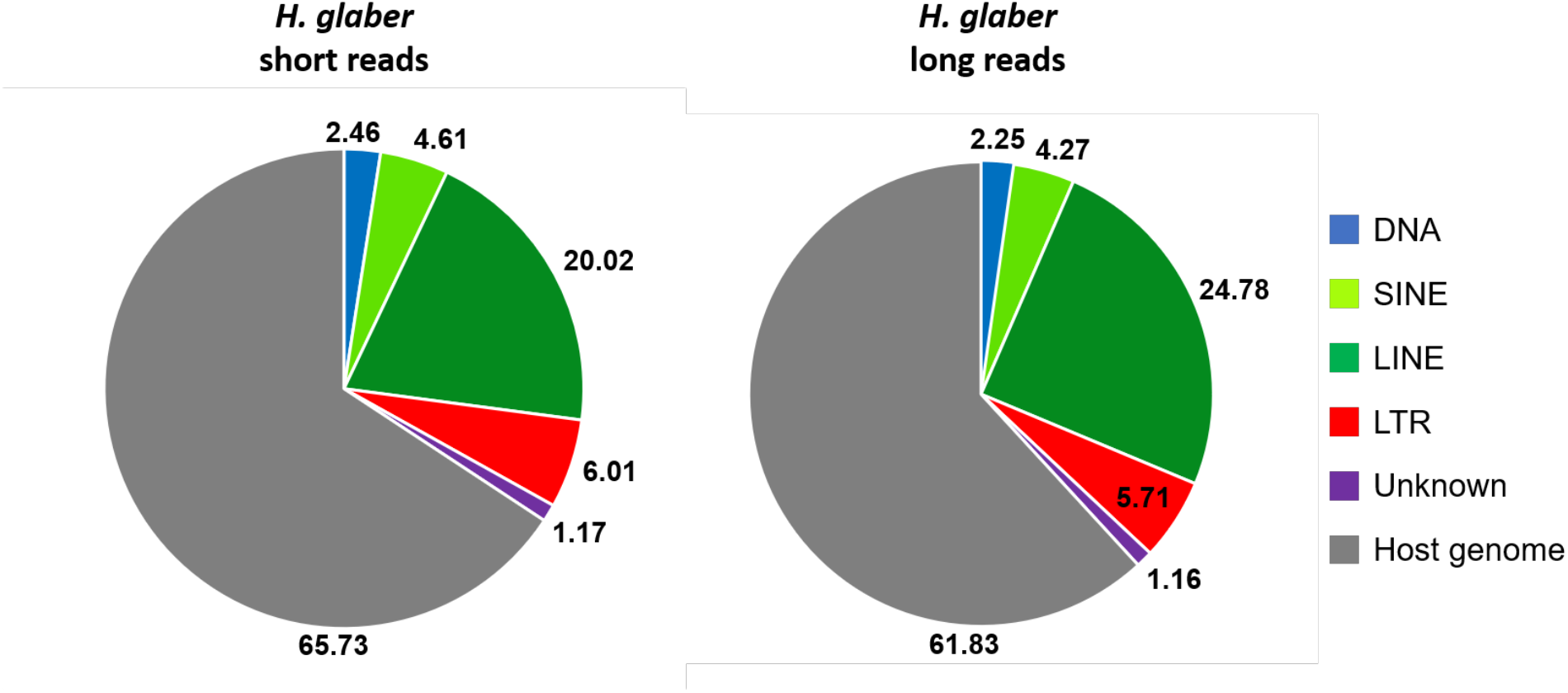
Transposable element content comparison between the two genome assemblies of the naked mole rat. The pie charts show the percentage of the main transposable element categories. The portion of the genome in grey comprises all the repetitive regions not annotated as transposable elements (e.g., tandem repeats and multi-copy gene families) as well as non-repetitive sequences. *H. glaber* short reads: HetGal_1.0, assembly size 2.6 Gb; *H. glaber* long reads: Heter_glaber.v1.7_hic_pac, assembly size 3 Gb.

**Figure 3.**
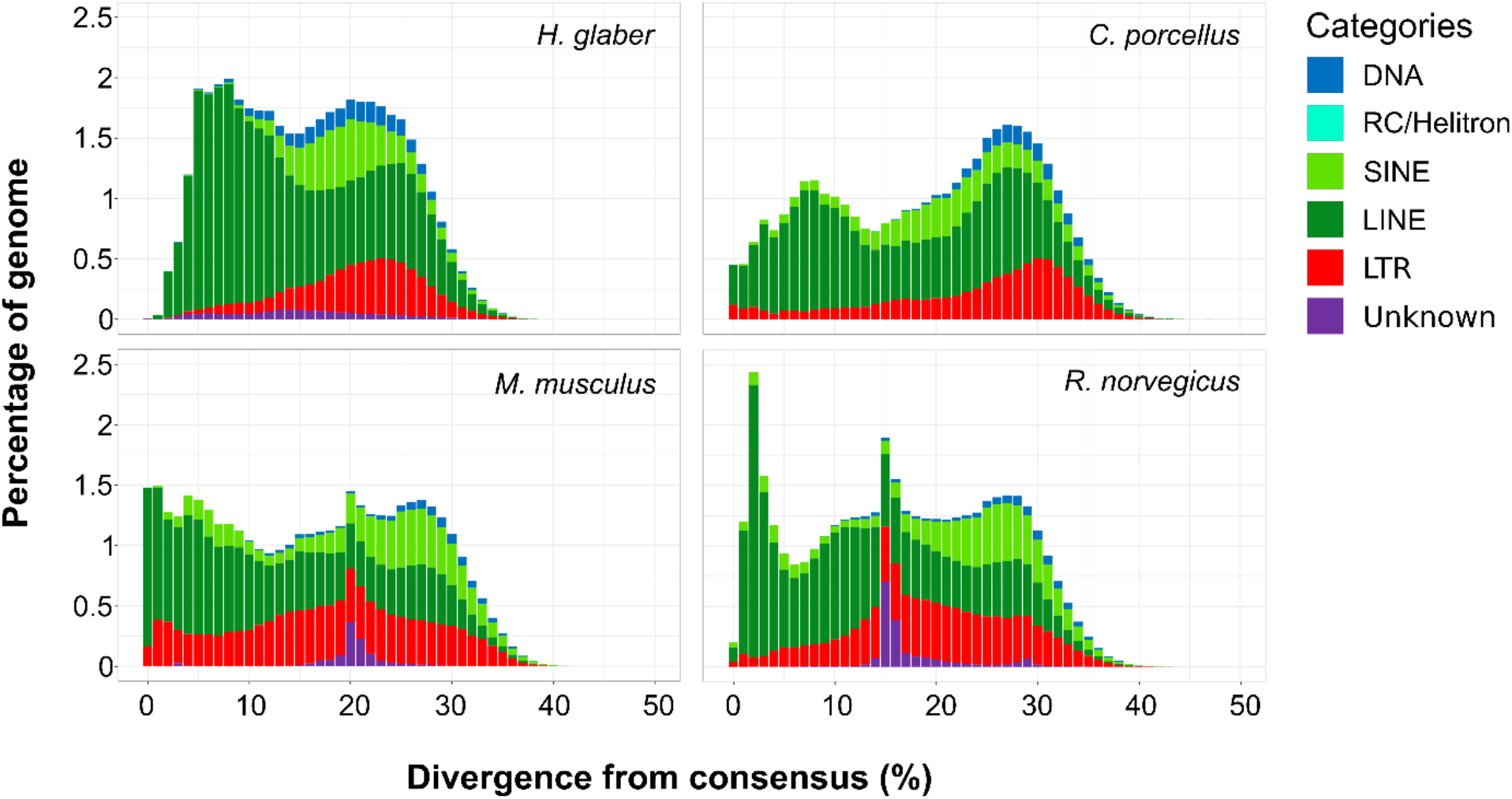
Transposable element landscapes of rodent species. The X-axis shows the genetic distance between the transposable element insertions and their consensus sequences, and the Y-axis shows the percentage of the genome occupied by transposable elements.

**Figure 4.**
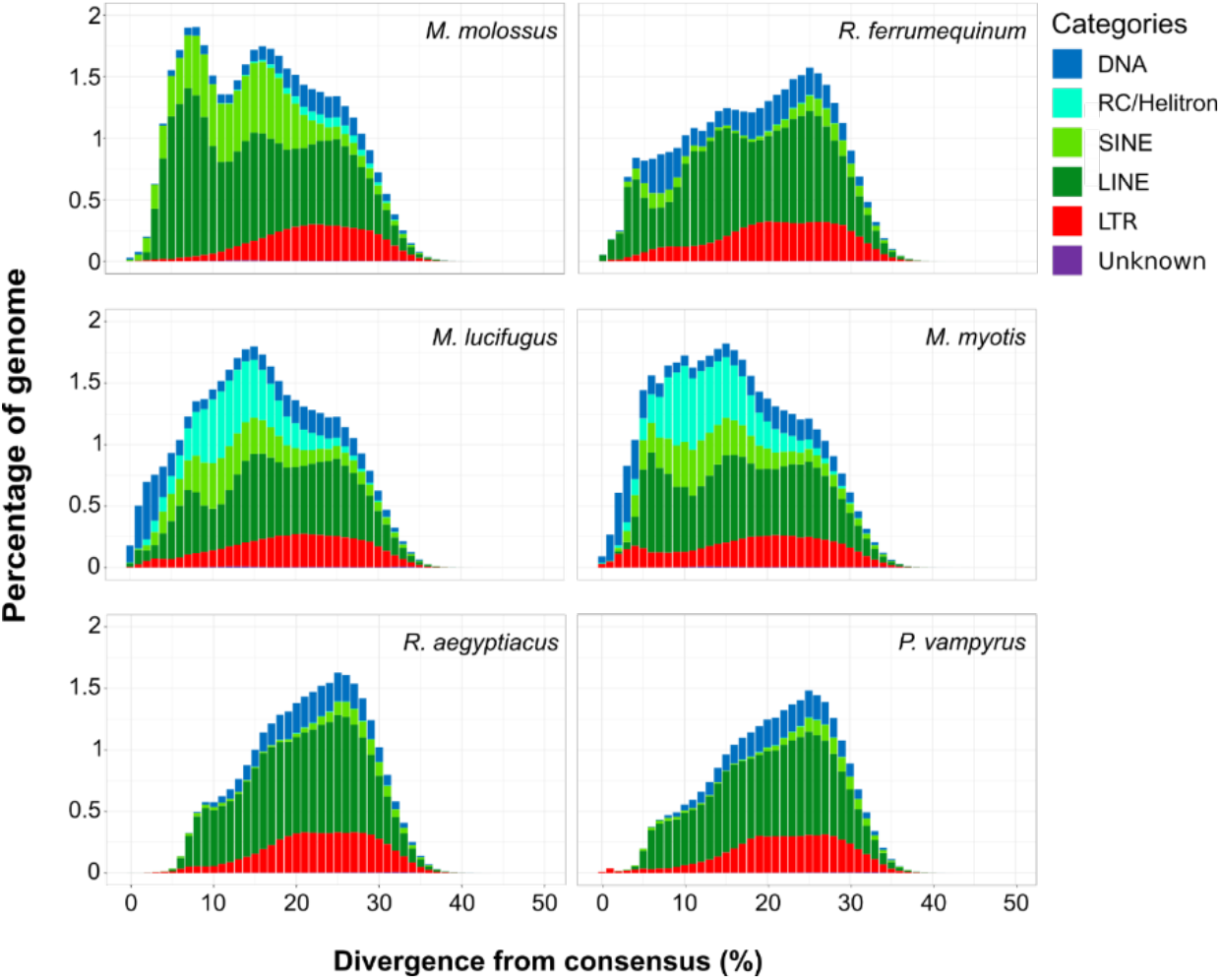
Transposable element landscapes of rodent species. The X-axis shows the genetic distance between the transposable element insertions and their consensus sequences, and the Y-axis shows the percentage of the genome occupied by transposable elements.

**Figure 5.**
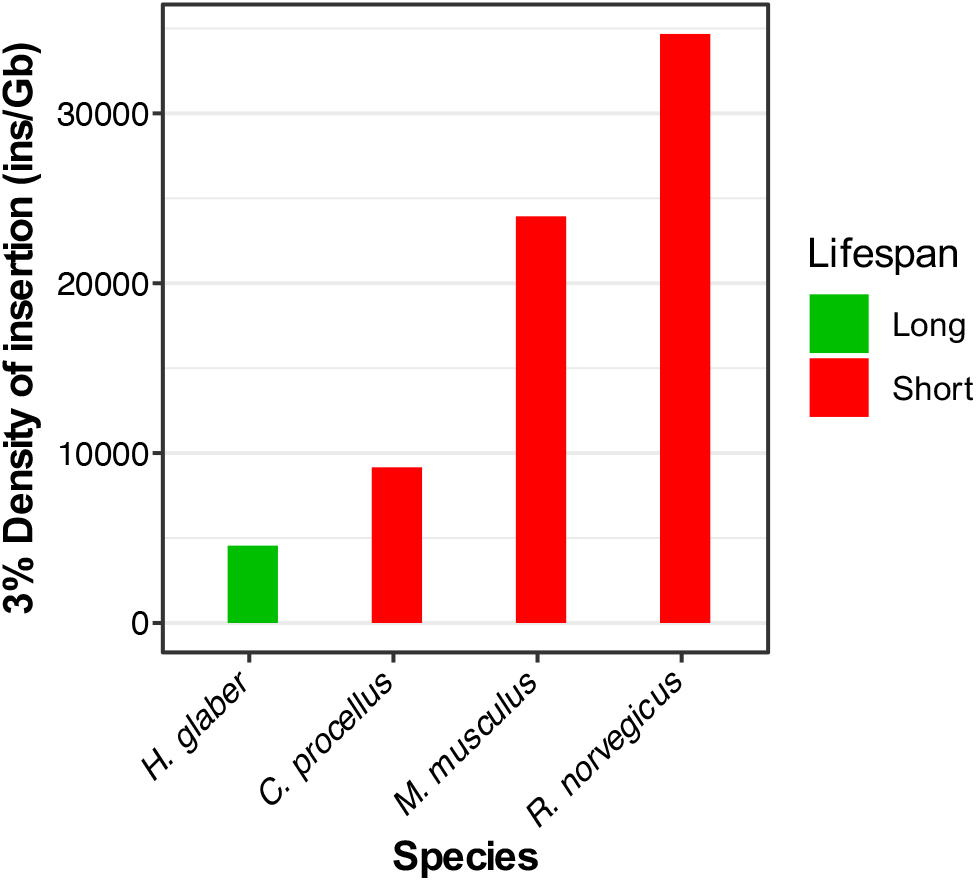
Density of insertion of recent non-LTR retrotransposons in rodents. The X-axis shows the species analyzed in decreasing order of longevity. The Y-axis shows the density of recent insertions per Gigabase (<3% divergence from consensus). Green indicates cancer-resistant and long-lived species. Red indicates cancer-prone and short-lived species. The black skull indicates the genomes in which intact LINE open reading frames were not found.

**Figure 6.**
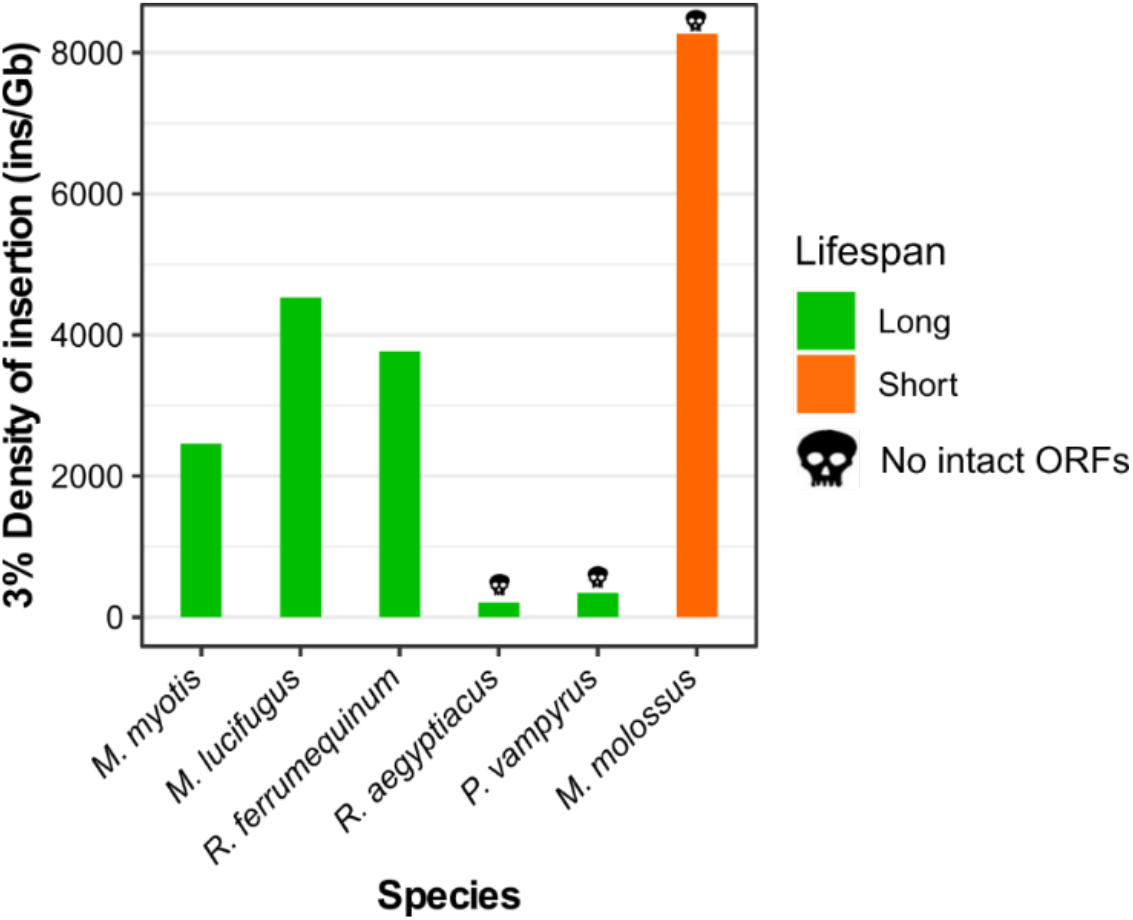
Density of insertion of recent non-LTR retrotransposons in bats. The X-axis shows the species analyzed in decreasing order of longevity. The Y-axis shows the density of recent insertions per Gigabase (<3% divergence from consensus). Green indicates cancer-resistant and long-lived species. Orange indicates cancer-resistant and short-lived species. The black skull indicates the genomes in which intact LINE open reading frames were not found.

### TE content in short- and long-read based assemblies of naked mole rat

The repetitive content of genome assemblies is tightly linked to the sequencing technology used. The long reads substantially help to reconstruct complex repetitive regions that remain hidden using exclusively short reads [1, 46]. For example, in the naked mole rat (*Heterocephalus glaber*), there are two male genome assemblies available: the first version (HetGla_1.0) sequenced using an Illumina platform and the latest version (Heter_glaber.v1.7_hic_pac) sequenced with PacBio. In the first analysis performed on HetGla_1.0 [47], the total TE content was 25% of the genome but such estimate is not available for Heter_glaber.v1.7_hic_pac [48]. Therefore, we produced a *de novo* repeat library based on the last assembly version and then annotated it. First, we ran RepeatModeler2 on Heter_glaber.v1.7_hic_pac to produce the *de novo* repeat library.

Then we used the newly produced repeat library to annotate the transposable elements in both assembly versions with RepeatMasker. The RepeatMasker annotation showed that the total transposable element content was 34.27% in HetGla_1.0 that increased to 38.17% in Heter_glaber.v1.7_hic_pac (**Figure 2**).

This latter percentage (38.17%) is more similar to the TEs content of the other rodents here analysed: 37.2% in *Cavia porcellus*, 41.88% in *Mus musculus*, and 41.13% in *Rattus norvegicus*. The TEs that showed the greatest increase between the two assemblies were the LINE retrotransposon (from 20.02% to 24.78%). In Heter_glaber.v1.7_hic_pac was also possible to observe a slight reduction of TEs classified as Unknowns (−0.01%), DNA transposons (−0.21%), LTR retrotransposons (−0.3 %) and SINEs (−0.34 %). On the whole, the mobilome of the naked mole rat showed a majority of retrotransposons and a minimal fraction of DNA transposons (less than 10% of the total TE content) in line with the TE content of other rodents and mammals.

### Transposable element analysis in rodents and bats

To investigate the differences in TE content between the rodent and bat species, we ran RepeatMasker on all their genome assemblies and the summary of the TE annotations are shown in **Table 1** and **Table 2**.

The genomes of the rodents and bats analysed here were mainly composed by class I retrotransposons (SINEs, LINEs, LTRs). The main difference between the two orders was that all six bats had a higher percentage of class II transposons (DNA transposons and Helitrons) than rodents, with the minimum percentage present in *M*. *molossus* (3.72%) and the maximum percentage in *M*. *myotis* and *M*. *lucifugus* (>11%; **Table 2**). Rodents, in general, showed a limited amount of class II transposons with the maximum abundance present in the naked mole rat (2.3 %). While the naked mole rat genome was the largest, it showed the lowest percentage of SINEs and LTR retrotransposons, and the highest percentage of LINEs among the analysed rodents (**Table 1**). Notably, *Myotis myotis* and *Myotis lucifugus* revealed an exceptional accumulation of Helitrons (6.93 and 6.18 % respectively) in comparison with the other bats that showed an abundance of Helitrons below 1%. The short-lived *M. molossus* showed contemporarily the largest genome size and the highest TE content (**Table 2**), in particular, the highest percentage of non-LTR retrotransposons (SINEs and LINEs).

To investigate how the TE content of rodents and bats changed over time and detect the most recently active transposable elements, we used the RepeatMasker annotation (see **Methods**) to generate a TE landscape for each of the species. The TE landscapes are a visualisation of the proportion of repeats in base pairs (Y-axis) at different levels of divergence (X-axis) calculated as a Kimura 2-p distance [49] between the insertions annotated and their respective consensus sequences (**Figure 3** and **Figure 4**). The TE landscapes in rodents were mostly dominated by retrotransposons and all of them showed an ancestral peak of accumulation between 20 and 30% of divergence (**Figure 3**). A small percentage of those ancient TEs was composed of relics of DNA transposons (blue in **Figure 3**). Notably, the cancer-resistant rodent naked mole rat showed the highest accumulation of retrotransposition dominated by LINEs (green) between 5 and 10% of divergence followed by a dramatic drop corresponding to the most recent history of this genome (**Figure 3**).

The three cancer-prone rodents showed an ongoing accumulation of LTR retrotransposons (red) and LINEs. *M. musculus* and *R. norvegicus* shared a peak of accumulation between 15 and 20% of divergence made of unknown interspersed repeats (purple). *R. norvegicus* was the only rodent that maintained a stable rate of retrotransposition of SINEs (light green).

The TE landscapes of the six bat species analysed (**Figure 4**) showed a higher inter-specific diversity in comparison with the four rodents described above (**Figure 3**). The most peculiar difference of the bat landscapes was the higher amount of class II transposons (**Table 2**). The two megabats belonging to the Pteropodidae family (*Rousettus aegyptiacus* and *Pteropus vampyrus*) presented the lowest TE accumulation (**Table 2**). Notably, *R. aegyptiacus, P. vampyrus* and *R. ferrumequinum* showed a shared peak of accumulation at 25% of divergence from consensus. The other three species (*M. myotis*, *M. lucifugus* and *M. molossus*) belong to the clade of Yangochiroptera evolved around 59 mya ago [50] and showed different landscapes. The two species belonging the *Myotis* genus shared their highest peak of accumulation at 15% of divergence, whereas *Molossus* had a second more recent species-specific peak at 8% of divergence. The two *Myotis* species showed similar landscapes likely due to the closer phylogenetical relationship. While *M. lucifugus* had a higher accumulation and diversity of TEs in its recent history (0-3% divergence), *Myotis myotis* had a longer history of accumulation of LTRs with a peak between 4 and 5% of divergence from consensus. In Chiroptera, the genus *Myotis* had the most pronounced accumulation of DNA transposons and Helitrons (light blue). In all the bats here considered, the non-LTR retrotransposons (LINEs and SINEs) showed a “*glaber*-like” dynamic with a decrease of retrotransposition in correspondence of their most recent evolutionary history (**Figure 4**).

### Activity of non-LTR retrotransposons in rodents and bats

We further explored the non-LTR retrotransposon accumulation (see **Methods**) by calculating the density of insertions (DI) from 0 to 3% of divergence. The density of insertion was estimated as the ratio between the number of non-LTR retrotransposon (SINEs and LINEs) insertions (**Figure 5** and **6**) and the genome size in Gigabases. The density of insertion is a sensitive parameter to compare the magnitude of the recent TE accumulation and its potential impact between species with different genome sizes [12]. When applying this measure in rodents (**Figure 5**), we found that the naked mole rat showed the lowest DI (4,145). All the cancer-prone species, that also have a shorter lifespan than the naked mole rat, showed a higher DI (**Figure 5**).

Then we calculated the density of insertions for the six bat species considered (**Figure 6**). All the bat genomes showed a lower DI than rodents with the exception of *Molossus molossus* that presented a DI of 8,475. The lowest DI (<1,000) was observed in *R. aegyptiacus* and *P. vampyrus*. The remaining species *M. myotis, M. lucifugus* and *R. ferrumequinum* had a DI ranging between 2000 and 5000. In general, the long-lived bats (indicated in blue in **Figure 6**) presented a lower DI compared to the short-lived *M. molossus* (**Figure 6**).

To investigate the possible presence of not only recently accumulated but active autonomous non-LTR retrotransposons we looked for the presence of LINE-related open reading frames (ORFs) with intact protein domains. By doing this, we found that all the analysed rodents genomes present intact LINE ORFs with the exclusion of the naked mole rat (**Figure 5**). In bats, intact LINE ORFs were found only in *M. myotis*, *M. lucifugus* and *R. ferrumequinum* (**Figure 6**).

## Discussion

Transposable element activity and accumulation can have manifold effects on genomes. Multiple studies have been linked transposable elements to ageing and the development of several diseases including cancer [22, 30, 31, 39, 51–54]. Here, we have studied the relationship between transposable elements and two different aspects of mammal life: longevity and cancer incidence. In rodents the short lifespan is associated with the presence of cancer [55]. On the other hand, bats are considered cancer-resistant species [56, 57] (**Figure 1**). The animals considered in this study are known to have different lifespans while sharing similar, small, body sizes (> 2 kg). *H. glaber* is the rodent with the longest lifespan known (31 years) and resistant to cancer while the other rodents show shorter lifespans of 12 (*C. porcellus*), 4 (*M. musculus*) and 3.8 (*R. norvegicus*) years. Chiroptera species, as far as it is currently known, are resistant to cancer (only a few confirmed cases have been reported [58, 59]) and show long lifespans for being small mammals [56] with a large variability (from 5.6 years of *M. molossus* to 37 years of *Myotis myotis*). Since TEs have been extensively linked to the development of cancer, we investigated the TE content of six bat species to see if it is possible to find shared features between long-lived bat and rodent species.

To analyse the transposable element content of these species, we relied on the use of high-quality genome assemblies based on long reads that better represent the actual genomic repetitive content with respect to assemblies based on short reads [1]. The use of long read-based genome assemblies is particularly important when analysing young TE insertions (as in this study) that are highly homogeneous and tend to be underrepresented in short read assemblies. On top of that, the combination of long read assemblies and custom TE libraries maximise the representation and annotation of the transposable element content as highlighted here for the genome of *H. glaber* (Figure 2). The first TE annotation of *H. glaber* without a custom TE library showed a repetitive content of about 25% [47] but the use of a proper TE library increased the content to ~34% (**Figure 2**) and, finally, the employment of long reads increased the total content to 38% (**Figure 2**, **Table 1)**.

We then proceeded with a more detailed analysis of the TE accumulation over time for rodents and bats to investigate possible differences in accumulation between short- and long-lived species and to find shared features between long-lived species. The main difference between short- and long-lived species of rodents is represented by a stark drop in non-LTR retrotransposon accumulation at recent times (0-5% divergence; **Figure 3**). Similarly, the long-lived species of bats show a drop in non-LTR retrotransposon accumulation at recent times (**Figure 4**) while presenting an overall accumulation of class II transposons (DNA transposons and Helitrons). Previous studies hypothesised that bats have a higher tolerance for the activity of transposable elements with alternative ways to dampen potential health issues due to this activity [41, 56]. Given our observation about the common drop of non-LTR retrotransposon accumulation in bats and naked mole rat, we add to the aforementioned hypothesis that it is the particular repression of the activity of non-LTR retrotransposon that enhances the resistance to cancer. The non-LTR retrotransposons are the most present types of TEs in rodents (**Figure 3**, **Table 1**) and the most extensively investigated by biomedical research given that they are the only active TEs in the human genome [11]. Since the TEs that may pose the highest health threat to organisms are the ones presently active, we compared the most recent accumulation of non-LTR retrotransposons between the long-lived and short-lived species considered in this study (**Figure 5**, **Figure 6**) using the density of recent insertions (DI). The naked mole rat showed the lowest DI of non-LTR retrotransposons followed by *C. porcellus* that has a lower cancer incidence than *M. musculus* and *R. norvegicus* [55] (**Figure 5**). Given the pattern of DI observed in rodents, the recent accumulation of non-LTRs appears to be related to the lifespan of these species (**Figure 1a**, **Figure 5**, **Figure 7a**). On top of that, the genome of the naked mole rat does not contain intact ORFs of LINE retrotransposons which is another clue for the absence of currently active non-LTR retrotransposons (**Figure 5**). The absence of intact LINE ORFs in the long read genome assembly confirms the same observation previously reported on the short read based genome assembly [60]. In bats, we observed that *M. molossus*, the bat species with the shortest lifespan, showed the highest DI, though no intact LINE ORFs were found in its genome. The high DI in *M. molossus* genome (**Figure 6**) is driven by a striking accumulation of non-autonomous SINEs. SINEs are about 300 bp long and do not present any protein coding sequence which make them unable to retrotranspose autonomously. Indeed, SINEs rely on the exploitation of the retrotransposition machinery provided by LINEs to move throughout the genome (**Figure 7**). Since *M. molossus* genome does not present any intact LINE ORFs, it is reasonable to consider both LINEs and SINEs to be inactive in this genome (**Figure 7b**).

**Figure 7.**
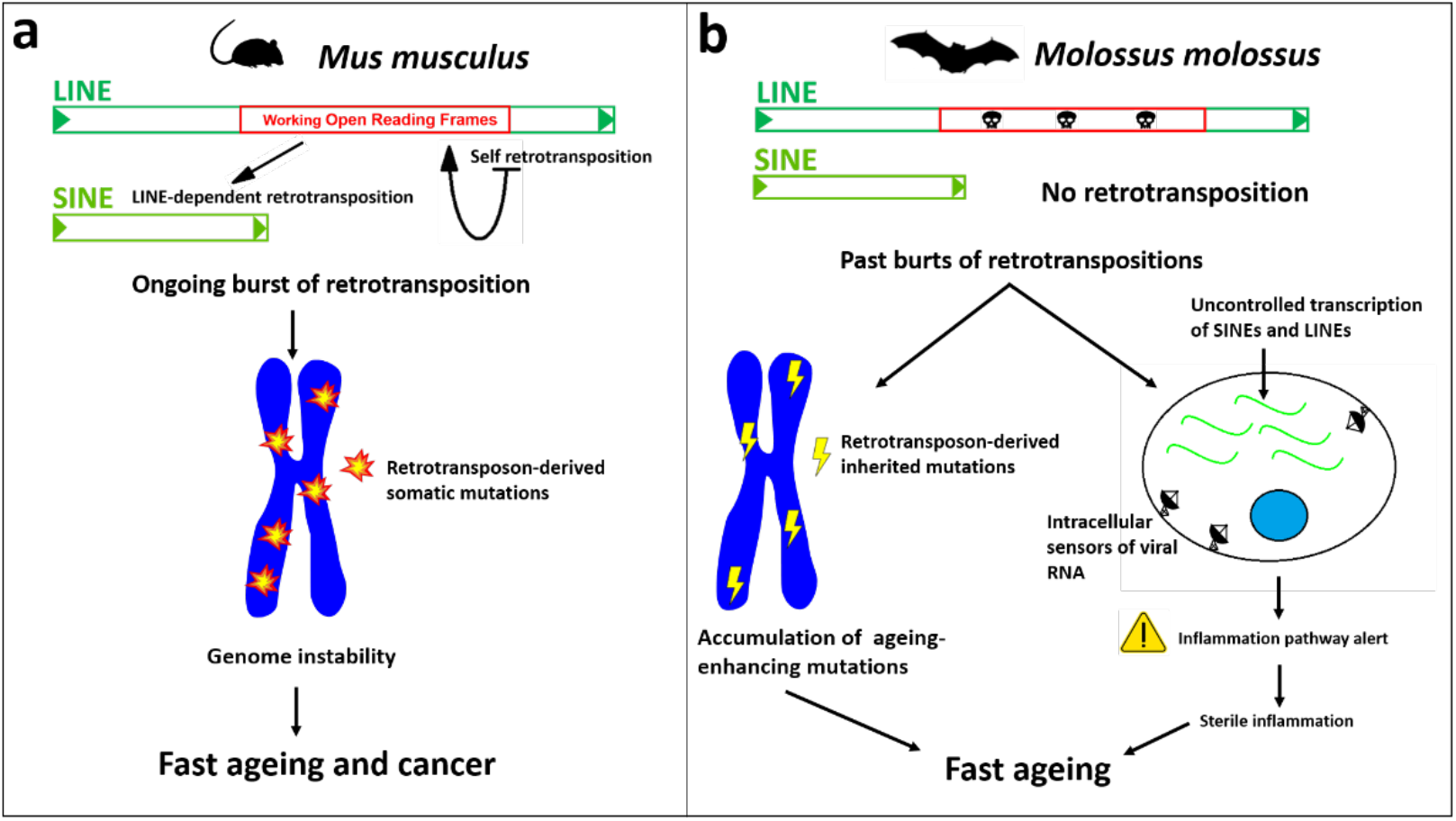
Hypothetical contribution of non-LTR retrotransposons to the fast ageing of rodents and bats. **a**) *Mus musculus* genome bears active non-LTR retrotransposons that destabilize the genome with continuous events of retrotransposition causing cancer and other diseases that reduce the mouse lifespan. **b**) *Molossus molossus* is both a cancer-resistant and short-lived species. The high density of non-LTR retrotransposons derives from a past accumulation of insertions (**Figure 4**) that may have resulted in the accumulation of ageing enhancing mutations and the continuous transcription of these retroelements may trigger cellular inflammation and cause a widespread sterile inflammation. In this scenario, the fitness of *M. molossus* is not impaired but its lifespan is reduced.

Theoretically, the absence of recent and active non-LTR retrotransposon insertions could make the genome more stable and reduce the frequency of cancer incidence. Nonetheless, the recently inserted SINEs and LINEs may still play a role in the physiology and ageing of *M. molossus*. A large number of very similar insertions across the chromosomes may cause structural polymorphisms (e.g., through non-allelic homologous recombination events) independently from the retrotransposition itself. For example, in the recent history of the human genome, LINE1-driven ectopic recombination events caused genomic rearrangements that can be responsible for diseases [14]. Moreover, it has been observed in the African killifishes that an accumulation of detrimental polymorphisms following a burst of TEs can undergo phases of relaxed selection were the mutations eventually promote age-associated diseases and reduced lifespan [61]. Among the killifishes, the turquoise killifish has a lifespan of 4-8 months (likely the shortest lifespan among vertebrates [61]) but despite the accumulation of mutations that leads to a fast ageing, the species is fully adapted to his ecological niche demonstrating that a rapid ageing is not necessarily linked to reduced fitness (**Figure 7b**). In addition to being considered genomic mutations, SINEs can also exert physiological effects by being transcribed. For example, in humans, SINEs of the Alu family have been found to be involved in the inflammatory pathway: upon viral infection, the interferon 1 mediates the overexpression of Alus which function as an amplifier of the immune response by stimulating the intracellular viral RNA sensor systems [62]. Despite, the involvement of SINEs in the human immune response, the same upregulation of Alus, or the excessive sensitivity of the viral RNA sensors, has been linked to autoimmune diseases [63–65]. Furthermore, as non-coding RNAs are emerging as essential components of neuroinflammation, a study has included Alu SINEs as possible promoters of neurodegenerative diseases [66]. More in general, the increased transcription of non-LTR retrotransposons (both LINEs and SINEs) in humans may contribute to the so-called “sterile inflammation” (**Figure 7b**), a phenomenon for which a chronic state of inflammation is triggered without the presence of any obvious pathogen and that is exacerbated with age (“inflammaging”) [67]. Therefore, it is possible that the great accumulation of SINEs in *M. molossus* trigger phenomena similar to the sterile inflammation and inflammaging that cause a shortening of its lifespan. For these reasons *M. molossus* may be an exceptional model for biogerontology.

## Conclusions

An increasing number of studies are finding a link between TEs, ageing, and cancer. As discussed in previous studies [39, 41], it is reasonable that genome instability derived by TEs could facilitate age-associated pathologies. In this work, we showed a relationship with retrotransposons activity and cancer incidence in rodents. The only cancer-resistant rodent species (naked mole rat) has a genome devoid of active LINEs and the lowest density of recently inserted non-LTR retroelements (SINEs and LINEs). The six analyzed species of bats are all cancer resistant, but they show widely different lifespans (from 37 years *Myotis myotis* to less than 6 years *Molossus molossus*). This particular taxon of mammals shows, together with a general cancer resistance, a drop of accumulation (and/or activity) of non-LTR retrotransposons in recent times similarly to the naked mole rat. The similar drop in non-LTR retrotransposon accumulation may indicate a possible involvement of retrotransposons in the incidence of cancer. The fast-ageing *M. molossus* has the highest density (DI) of non-LTR retroelements among bats (**Figure 6**) driven by the accumulation of SINEs. Despite the higher accumulation of SINEs in *M. molossus*, the absence of intact LINEs (that mobilise SINEs) makes SINEs inactive and not a source of new somatic mutations through their retrotransposition. This genomic feature makes *M. molossus* theoretically as cancer resistant as the other bat species. In humans and other mammals non-LTR retrotransposons are also part of the inflammation pathway [REF]. The high density of SINEs observed in *M. molossus* may promote ageing trough their high mutation load and their pervasive transcription may participate to the processes similar to the sterile inflammation and inflammaging. In conclusion, the non-LTR retrotransposon content may act as promoting factor, more than as simple biomarker, in cancer evolution and other age-associated conditions.

## Methods

### Samples

In order to avoid biases related to potential underestimations of the TE content, in this study we chose species of mammals from which high-quality genome assemblies were available. The 10 species taken in to account and the accession numbers of their assemblies are listed here: *Mus musculus* GCA_000001635.8 [ Genome Reference Consortium mouse reference 38], *Rattus norvegicus* GCA_000001895.4 and [68], *Cavia porcellus* GCA_000151735.1[69], *Heterocephalus glaber* GCA_014060925.1 [48], *Rousettus aegyptiacus* GCA_014176215.1 [70], *Rhinolophus ferrumequinum* GCA_014108255.1 [70], *Molossus molossus* GCA_014108415.1 [70] *Pteropus vampyrus* GCA_000151845.1 [69], *Myotis myotis* GCA_014108235.1 [70], *Myotis lucifugus* GCA_000147115.1 [69]. The species selected are from Rodentia and Chiroptera orders. These two orders have been separated in the Mesozoic era 90 mya ago [71]. The large evolutionary distance among these two orders allowed us to find genomic features potentially conserved in all mammals.

Body mass is generally positively related to the lifespan [45] therefore, in our study, we chose to analyse genomes belonging to small-sized mammals (between 12 g and 1.1 kg). The comparisons were carried out between species belonging to the same taxonomical order to maximize their shared evolutionary history, physiological features and life history traits, then the species were also compared on the basis of their lifespans (**Figure 1**). The genome assemblies analysed are all based on Sanger or PacBio technologies to maximize their assembled repetitive content [72–75]. The short (GCA_000230445.1 [47]) and long read based genome assemblies of *H. glaber* were compared in order to estimate the differences in TE content due to technological biases.

### Repetitive element libraries and annotation

To analyze the content of transposable elements in a genome, the genome assemblies must be annotated with proper libraries of TE consensus sequences. A consensus represents the reference sequence of a specific subfamily of transposable elements. The repeat annotation softwares (like RepeatMasker) make use of such library of consensus sequences to find instances (or insertions) of each repeat in the genome assemblies. From the alignments between the insertions found in the genome assembly and their consensus sequences, it is possible to estimate the age of such insertions on the basis of the genetic distance given by the alignments: the higher the distance, the higher the age of the insertions.

All the species analysed have a species-specific repeat library already available but *Heterocephalus glaber*. Therefore, we made a *de novo* repeat library of *H. glaber* using RepeatModeler2 with the option -LTRstruct [76] The resulting consensus sequences were merged with the ancient mammalian TE families (LINE2 and MIR SINEs) downloaded from RepBase (https://www.girinst.org/repbase). All the genome assemblies of rodents (but *H. glbaer*) were masked using RepeatMasker (v. 4.1.0) and the Rodentia-specific library (Repbase release 20181026). Similarly, the bat genome assemblies were masked using the Chiroptera-specific library from Repbase merged with the curated TE libraries produced by Jebb et al. [70]. All RepeatMasker annotations were performed using the options -a -xsmall -gccalc -excln. The TE annotations were then visualized through barplots (landscapes) that show in the X-axis the genetic distances from the consensus sequences (calculated as Kimura 2-p distance [49]) and in Y-axis the percentage of the genome occupied by each TE category. The barplots were made using the ggplot2 R package [77]. For each species, the genome assembly size and the percentage of each TE categories listed in **Table 1**, **Table 2** and **Figure 2** were retrieved from the table files produced by RepeatMasker.

### Density of young non-LTR retrotransposons

To evaluate the density of recent non-LTR retrotransposon insertions (LINEs and SINEs), we selected all the non-LTR retrotransposon hits found by RepeatMasker with a divergence from consensus lower than 3%. Then we calculated the density of insertion (DI) [12] for each species as the ratio between the number of recent insertions and the corresponding assembly size expressed in Gigabases:

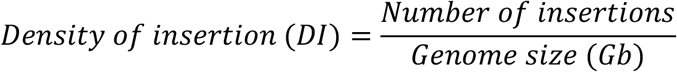

### Intact ORF detection

To check the presence of intact LINE retrotransposons (therefore potentially active) we looked for complete open reading frames (ORFs) that encode for the enzymatic machinery used by LINEs to retrotranspose themselves and the non-autonomous SINEs. In each species the sequences of LINEs were with BEDTools getfasta using the LINE coordinates annotated by RepeatMasker. A fasta file for each species was produced and used as input for the R script orfCheker.R (https://github.com/jamesdgalbraith/OrthologueRegions/blob/master/orfChecker.R) to find intact LINE ORFs. The script considers a LINE ORF to be intact if it contains both complete reverse transcriptase and endonuclease domains.

## Acknowledgments

This study has been funded by MAPS Department funds BIRD2021 - prot. BIRD213010, University of Padova.

